# An alternative nisin A resistance mechanism affects virulence in *Staphylococcus aureus*

**DOI:** 10.1101/716191

**Authors:** Miki Kawada-Matsuo, Atsuko Watanabe, Kaoru Arii, Yuichi Oogai, Kazuyuki Noguchi, Shouichi Miyawaki, Tetsuya Hayashi, Hitoshi Komatsuzawa

**Affiliations:** Department of Oral Microbiology, Kagoshima University Graduate School of Medical and Dental Sciences, Kagoshima, Japan; Department of Orthodontics and Dentofacial Orthopedics, Kagoshima University Graduate School of Medical and Dental Sciences, Kagoshima, Japan; Department of Periodontology, Kagoshima University Graduate School of Medical and Dental Sciences, Kagoshima, Japan; Department of Bacteriology, Faculty of Medical Sciences, Kyushu University, Fukuoka, Japan; Department of Bacteriology, Hiroshima University Graduate School of Biomedical and Health Sciences, Hiroshima, Japan

**Author notes:** These authors contributed equally to this work (Miki Kawada-Matsuo, Atsuko Watanabe). **Abbreviations:** hBD3, human beta-defensin-3; MIC, minimum inhibitory concentration; PBS, phosphate buffered saline; PSM, phenol soluble modulin; TCS, two-component system; TSA, tryptic soy agar; TSB, tryptic soy broth.

## Abstract

Nisin A is a bacteriocin produced by *Lactococcus lactis* and is widely used as a food preservative. *Staphylococcus aureus* has the BraRS-VraDE system providing resistance against low concentrations of nisin A. The BraRS is one of a two-component system that senses nisin A by BraS and finally induces the expression of ABC transporter VraDE by phosphorylated BraR. Previously, we isolated a highly nisin A resistant strain with increased VraDE expression due to a mutation of *braRS*. In this study, we isolated a BraRS-VraDE-independent, nisin A resistant mutant from *S. aureus* MW2. These mutants, designated SAN2 (*<u>S</u>. <u>a</u>ureus* <u>n</u>isin resistant) and SAN469, had a mutation in *pmtR* (MW1875) which encodes a transcriptional regulator responsible for the expression of the *pmtA-D* operon. As a result, this mutant exhibited a high level of constitutive production of PmtA-D, a transporter responsible for the export of phenol soluble modulin (PSM). We also obtained two *pmtA-D* overexpressing, nisin A resistant mutants which contained a point mutation in *pmtR* from other *S. aureus* strains.

Characterization of the mutants revealed that they have a decreased susceptibility to human beta defensin-3 and LL37, which are innate immune factors. Additionally, these mutants showed higher hemolytic activity than the MW2 original strain. Furthermore, in a mouse bacteremia model, the SAN2 strain exhibited a lower survival rate than the MW2 original strain. These results indicate that the over expression of *pmtA-D* due to the *pmtR* mutation is an alternative nisin A resistance, which also affects virulence in *S. aureus*.

**Author Summary:** Recently, the emergence of antibiotic resistant bacteria such as MRSA, MDRP and CRE have brought serious problems for chemotherapy in the world. In addition, many antibacterial agents such as disinfectants and food additives are widely used. Therefore, it raises the possibility that bacteria are becoming resistant to all antibacterial agents. In this study, we investigated whether *S. aureus* become resistant against nisin A, one of the food additives. Finally, we isolated nisin A highly resistant *S. aureus* strains. Among these strains, we identified that one strain designated as SAN2 showed nisin A resistance by the overproduction of Pmts which were involved in the secretion of virulence factors called PSMs. We identified a mutation of *pmtR* gene encoding a regulator for *pmt* genes. SAN2 strain showed the decreased susceptibility to human antimicrobial peptides and the increased hemolytic activity. Finally, SAN2 showed higher lethal activity in mouse bacteremia model. Our study provides new insights into that *S. aureus* may cause resistance against various antibacterial food additives, together with the altering the virulence.

## Introduction

Bacteriocins are defined as antimicrobial peptides produced by bacteria. Many bacteriocins have been reported in a wide range of bacterial species including Gram-positive and gram-negative bacteria (1–4). Bacteriocins are mainly divided into two groups, lantibiotics and nonlantibiotics. Lantibiotics are antimicrobial peptides that contain unusual amino acids, the lanthionines (5, 6). Nisin A is one of the lantibiotics produced by *Lactococcus lactis* (7). The target of nisin A is lipid II, which is involved in cell wall biosynthesis (8, 9). The binding of nisin A to lipid II inhibits cell wall biosynthesis and eventually causes the formation of pores or disturbances in bacterial membranes. Generally, bacteriocin has a strong antibacterial activity against closely related species. However, nisin A has a broad range of antibacterial activities mainly against gram-positive bacteria, including species of *Streptococcus, Staphylococcus* and *Clostridium* (10–15). Due to its broad range activity, nisin A is widely used as a food additive throughout the world for the prevention of food-borne poisoning (10, 16, 17). Additionally, the use of bacteriocins, including nisin A, as clinical antibacterial agents has been investigated (10, 13, 18).

*S. aureus* is a commensal bacterium in human; generally localizes in the nasal cavity, skin and intestine; and sometimes causes opportunistic infections such as suppurative diseases, pneumonia and sepsis (19–21). Additionally, *S. aureus* causes food-borne poisoning because this organism produces several heat-stable enterotoxins (20). Since some foods contain nisin A as a preservative, *S. aureus* is sometimes exposed to nisin A (10, 22). Previously, we and other groups have demonstrated that a two-component system (TCS) named BraRS is responsible for resistance to nisin A (23, 24). When *S. aureus* cells are exposed to nisin A, BraS senses nisin A, and phosphorylation of BraR then occurs. Phosphorylated BraR binds upstream of *vraDE* which encodes an ABC transporter responsible for nisin A resistance. However, a high concentration of nisin A is still effective against *S. aureus* cells (14). Previously, to determine whether nisin A treatment induces higher nisin A resistance, we tried to isolate nisin A mutants that showed increased levels of nisin A resistance by the incubation with sub-minimum inhibitory concentration (MIC) of nisin A and finally obtained three mutants that constitutively expressed high level of VraDE (25). We found point mutations in the promoter region of *braRS,* the *braR* or *braS* coding region in each mutant. In this study, to explore potential nisin A resistance mechanisms independent of the BraRS-VraDE system, we tried to isolate *S. aureus* strains highly resistant to nisin A by exposing *S. aureus* to nisin A, and eventually obtained BraRS-VraDE independent nisin A resistant mutants. Our analysis of these mutants demonstrates that they utilize an alternative nisin A resistance mechanism and the mutations also affect the virulence of *S. aureus*.

## Results

### Isolation of *S. aureus* strains highly resistant to nisin A

Previously, we obtained three types of *S. aureus* strains highly resistant to nisin A, which showed increased levels of *vraDE* expression (25). We further tried to isolate strains highly resistant to nisin A. From two independent experiments, we obtained two mutants that showed no increased expression of *vraDE*. We designated these mutants SAN2 and SAN469. The MICs of SAN2 and SAN469 against nisin A were 2,048 µg/ml, while the MIC of the MW2 original strain was 512 µg/ml.

We also evaluated the MICs of these mutants against bacitracin, gallidermin and nukacin ISK-1, which also inhibit the lipid cycle for cell wall biosynthesis (Table 3). There was no difference in MICs for these three agents between the mutants and the MW2 original strain.

### Analysis of gene expression in MW2 and SAN2 by microarray analysis

To identify the factors responsible for high resistance to nisin A, we investigated the expression of all genes located on the chromosome of MW2. As shown in Table 4, the expression of MW1875 to MW1871 was significantly increased in the SAN2 strain, with more than 30-fold higher expression levels than that of the MW2 original strain in the absence of nisin A. MW1875-MW1871 were previously associated with phenol-soluble modulin (PSM) transport and designated PmtR (MW1875) and PmtA - PmtD (MW1874-71) (30). It was reported that *pmtR* encodes the transcriptional regulator (PmtR), and PmtA-D constitute a PSM transporter. In strain SAN2, the expression levels of several other genes were also increased, but too much lower extents (3- to 5-fold greater than the MW2 levels). We also confirmed the increased expression of MW1875 to MW1871 in SAN469 by quantitative PCR with the method described in Materials and Methods.

### Effects of the inactivation of MW1875 or MW1874 on the nisin A susceptibility

From microarray analysis, we thought MW1875-MW1871 were associated with high resistance to nisin A. Therefore, we constructed corresponding inactivation mutants to examine whether these genes truly contributed to high nisin A resistance. Since *pmtR* and *pmtA-D* (MW1875-1871) formed an operon, we constructed two insertional mutants (MM2278 and MM2153), in which pYT1 was integrated in MW1875 or MW1874 of strain SAN2, and one insertional mutant (MM2202) with an insertion in MW1875 of strain MW2. The two SAN2 mutants showed the same MIC against nisin A as that of MW2. Similarly, the MW2 mutant (MM2202) retained the same MIC as that of MW2 (Table 5). A complement strain MM2279, in which the *pmtRABCD* operon (MW1875-1871) was expressed by pCL15 in MM2278, also showed a similar MIC as that of SAN2.

### DNA sequences of the MW1875 to 1871 (*pmtR* and *pmtA* to *D*) regions

The DNA sequences of the MW1875-1871 regions in MW2, SAN2 and SAN469 were determined. In SAN2, only one mutation was detected in MW1875 (Fig. 1). This mutation induced an alanine (Ala) at 43^rd^ amino acid of MW1875 to aspartic acid (Asp) in SAN2 strains. In SAN469, the fifth amino acid position in MW1875 was mutated into a stop codon (Fig. 1).

**Fig. 1.**
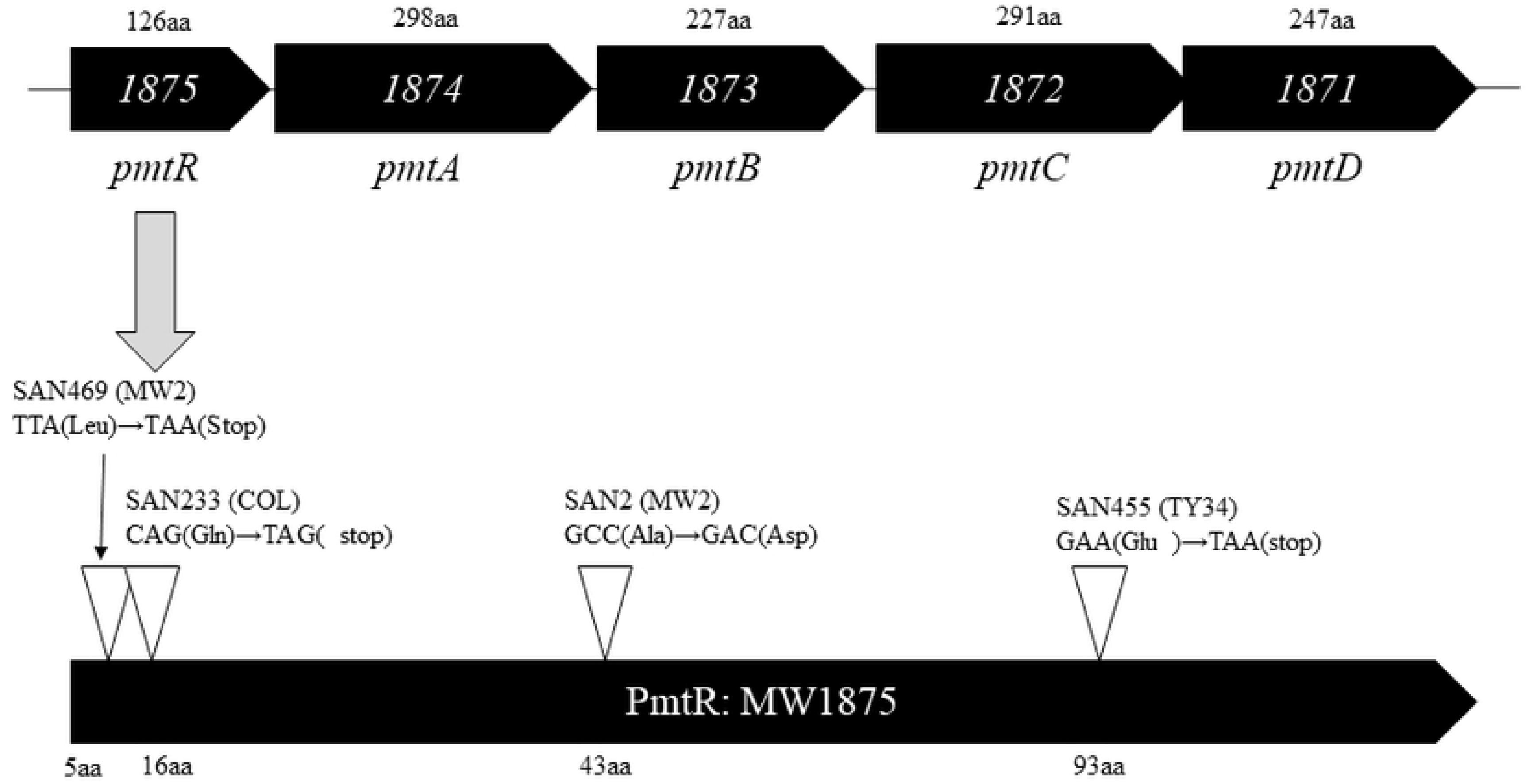
ORF map of the *pmt* region (MW1875-1871) and mutation sites in *ptmR* of isolated mutants. The mutation sites in the ptmR region are indicated by white arrows.

### Isolation of strains highly resistant to nisin A from other *S. aureus* strains

To determine whether similar *pmtR* mutants were able to be obtained from other strains by the exposure to nisin A, we isolated nisin A resistant mutants from *S. aureus* COL and TY34. From two independent experiments in each strain, we obtained two mutants with increased expression of *pmtR*A*BCD*, one from each strain. We designated the mutants SAN233 (from COL) and SAN455 (from TY34). DNA sequencing analysis identified point mutations at the 16^th^ and 93^rd^ amino acids of MW1875 (PmtR) in SAN233 and SAN455, respectively, while there was no mutation in the *pmtA*-*D* genes. These two mutations introduced stop codons within the *pmtR* gene. Therefore, SAN233 and SAN455 did not express full-length PmtR.

### Expression of *PmtR* (MW1875), *PmtA* (MW1874) and *vraD* in the MW2 and its mutants

We investigated the expression of *pmtR*, *pmtA* and *vraD* by quantitative PCR. In SAN2, the expression of *pmtR* and *pmtA* was significantly increased compared to that of the MW2 in the absence of nisin A (Fig. 2A). We observed the same result by immunoblotting analysis (Fig. 2B). In addition, the expression patterns of PmtR and PmtA in the absence of nisin A were quite similar to those in the presence of nisin A because the expression of *pmtR* and *pmtA* was not induced by nisin A (supplemental Fig. 2). Additionally, we investigated the expression of *vraD* and found that SAN2 showed no increase in *vraD* expression at a low concentration of nisin A (32 µg/ml), while MW2 showed increased expression of *vraD* (Fig. 2C). However, the inactivation of *pmtR*A*BCD* in SAN2 resulted in an increaseed *vraD* expression in the presence of nisin A.

**Fig. 2.**
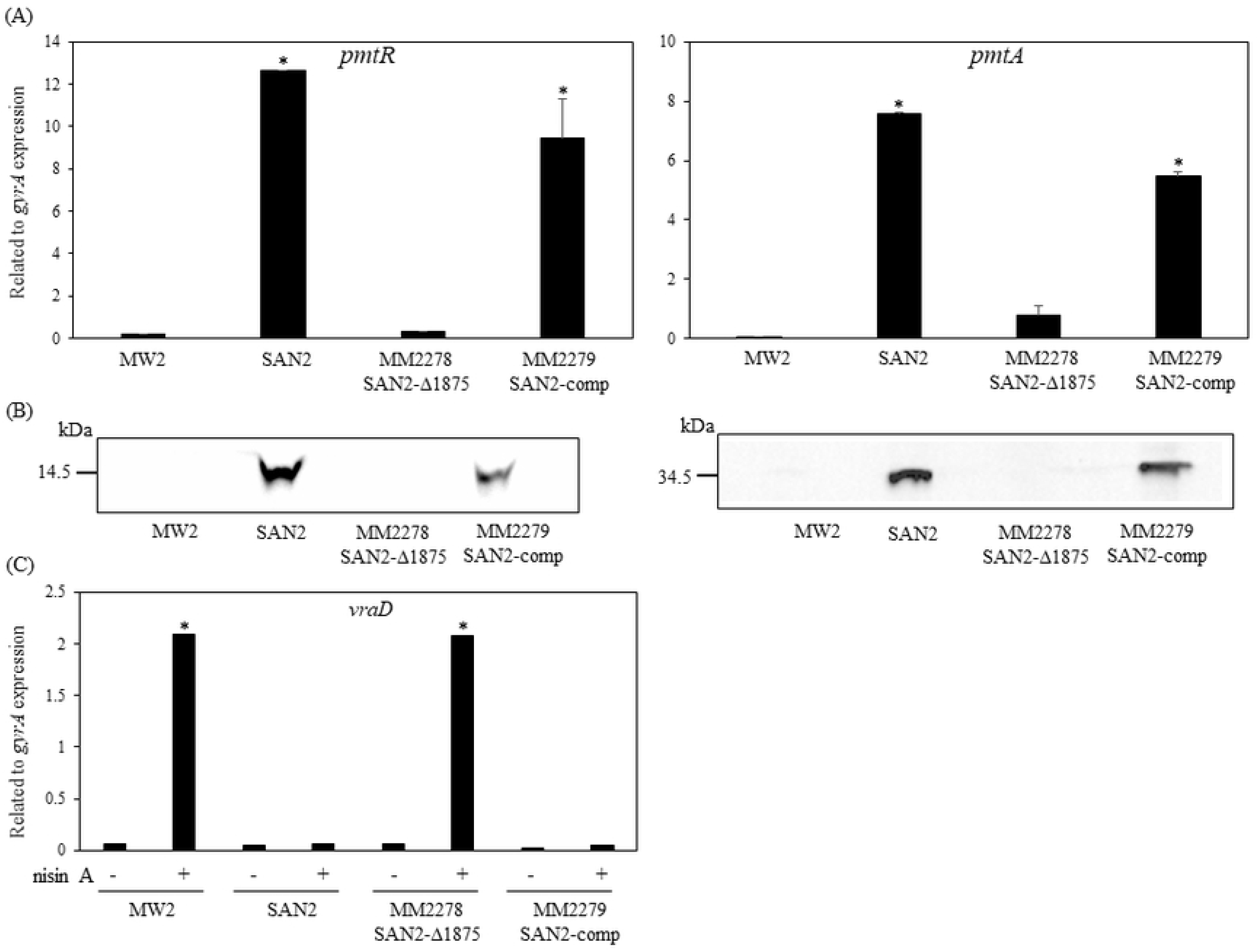
Expression of PmtR, PmtA and VraD in the mutants. (A) Expression of PmtR and PmtA in MW2, SAN2, and SAN2 with inactivated *pmtR-pmtD* and its complement strain were used for immunoblotting analysis. Exponential phase cells that reached OD_660_=0.5 were collected. Immunoblotting analysis using each specific antibody was performed as described above. \**p* <0.001, as determined by Dunnett’s post hoc tests compared to untreated MW2. (B) Expression levels of *pmtR* and *pmtA* in *S. aureus* strains were evaluated by quantitative PCR using specific primers. Exponential phase cells that reached OD_660_=0.5 were collected. Quantitative PCR using specific primers was performed as described above. (C) Expression of *vraD* in *S. aureus* strains in the presence or absence of 1/16 MIC nisin A was evaluated by quantitative PCR using specific primers. \**p* <0.005, as determined by Dunnett’s post hoc tests compared to untreated MW2.

Then, we investigate the *vraD* expression in the presence of various concentrations of nisin A (Fig. 3A) In MW2, the expression of *vraD* was induced at concentrations above 1/32 MIC nisin A (16 µg/ml), while in SAN2, it was induced above 1/2 MIC nisin A (1024 µg/ml). We further investigated the induction of *vraD* expression by bacitracin (1 MIC=64 µg/ml). Since the MIC of bacitracin was the same in the MW2 and SAN2 strains, we analyzed the effect of bacitracin at a range from 1/64 MIC to 1 MIC. The *vraD* expression in both strains was similarly induced by bacitracin at each concentration (Fig. 3B).

**Fig. 3.**
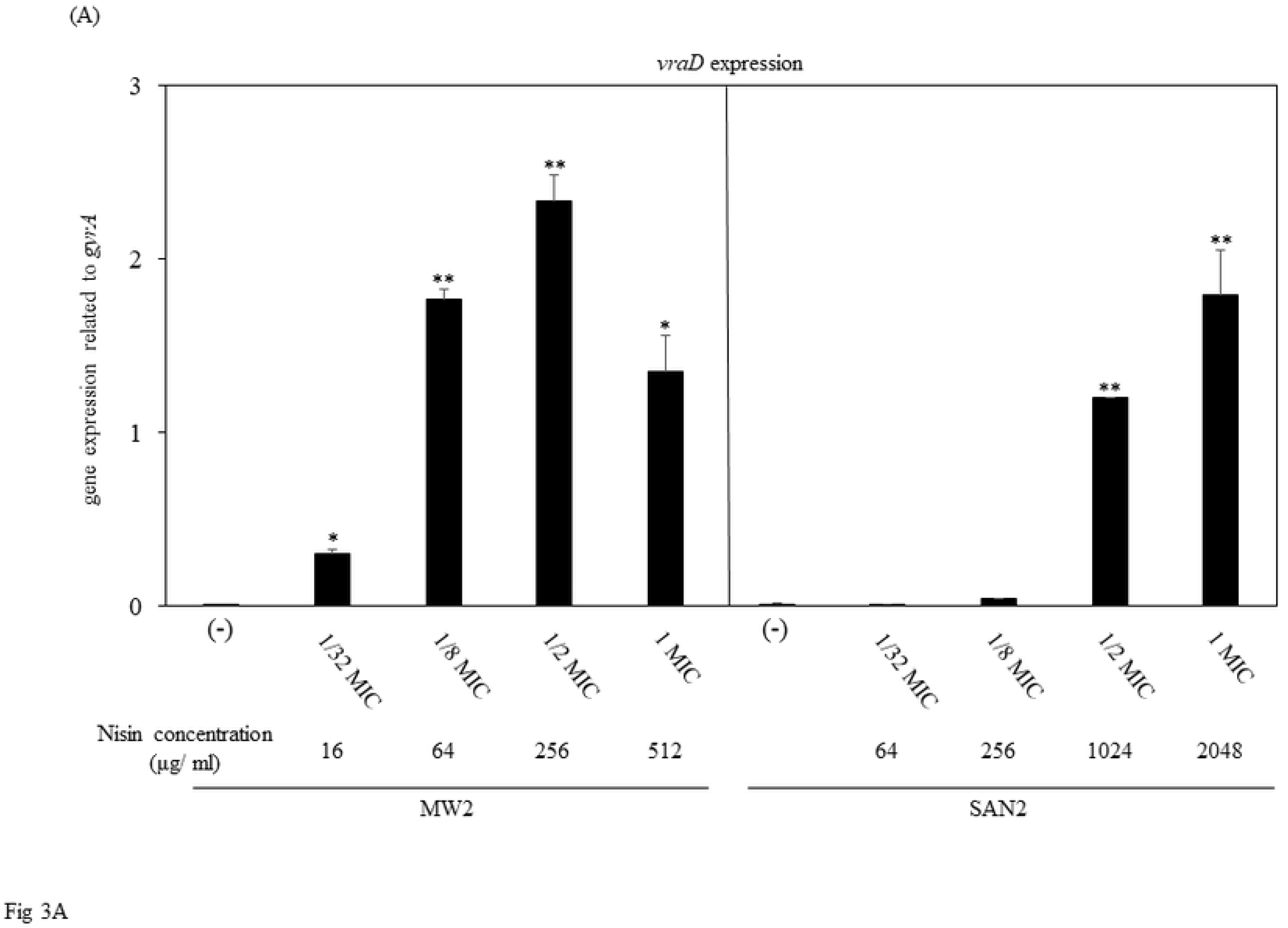
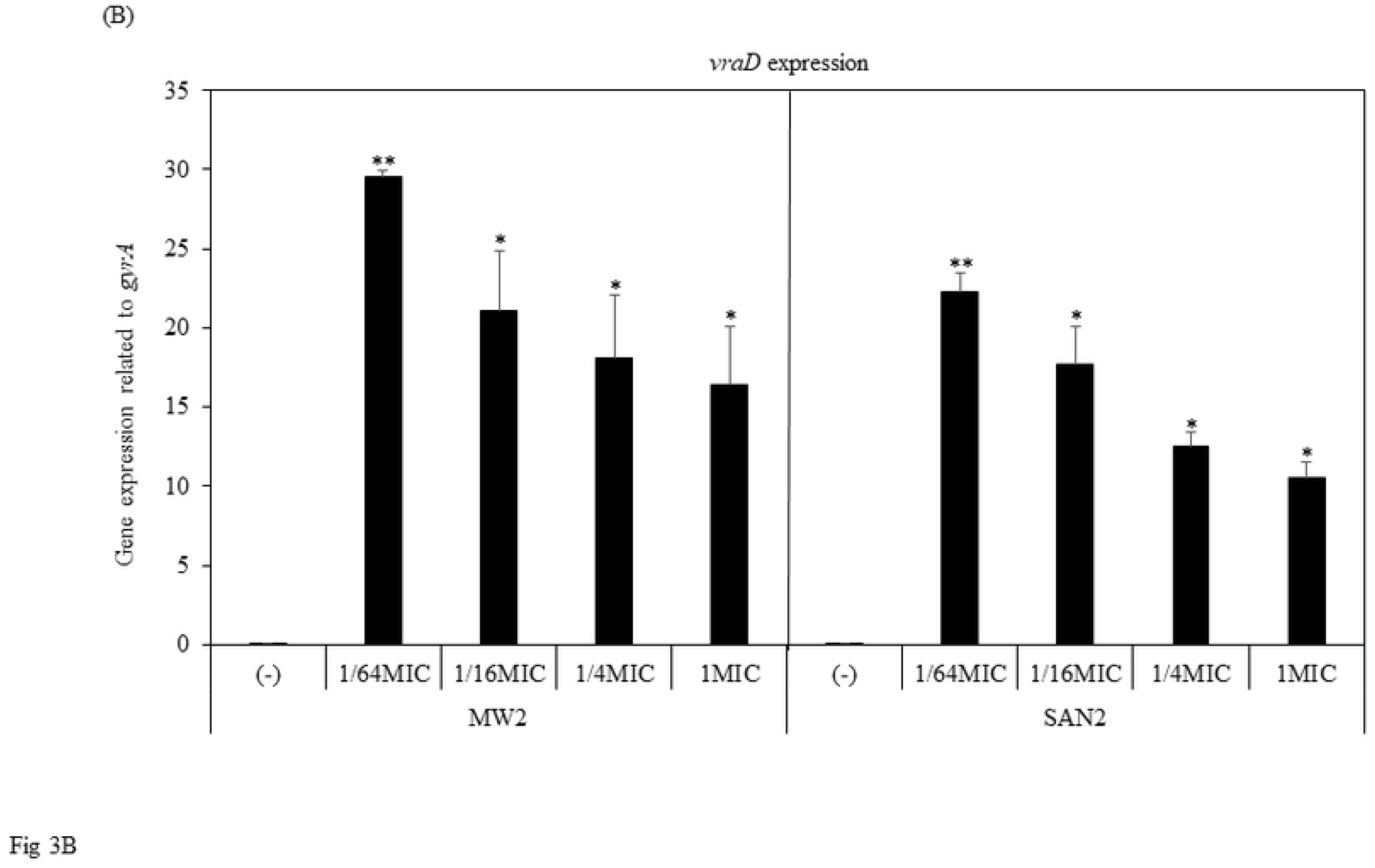
Expression of vraD in MW2 and SAN2 with various concentrations of nisin. **A.** Induction of *vraD* expression in *S. aureus* MW2 and SAN2 by addition of various concentrations of nisin A (A) or bacitracin (B) was evaluated by quantitative PCR using specific primers. Exponential phase cells that reached OD_660_=0.5 were exposed to various concentrations of nisin A or bacitracin. After incubation for 15 min, the bacterial cells were collected. RNA extraction, cDNA synthesis and quantitative PCR were performed as described above. \**p* <0.05, ** *p* <0.01 as determined by Dunnett’s post hoc tests compared to untreated MW2 or SAN2.

### Binding of the wild-type and mutated PmtR protein to the upstream region of MW1875

Fig. 4A shows the previously reported promoter region of *pmtR*, the transcriptional start site and the binding region of PmtR (31). The results of our EMSA revealed that MW2-rPmtR bound the upstream region of *pmtR* (*pmtR*-F), while SAN2-rPmtR did not (Fig. 4B). This binding of MW2-rPmtR was inhibited by the addition of an excess amount of unlabeled DNA fragment.

**Fig. 4.**
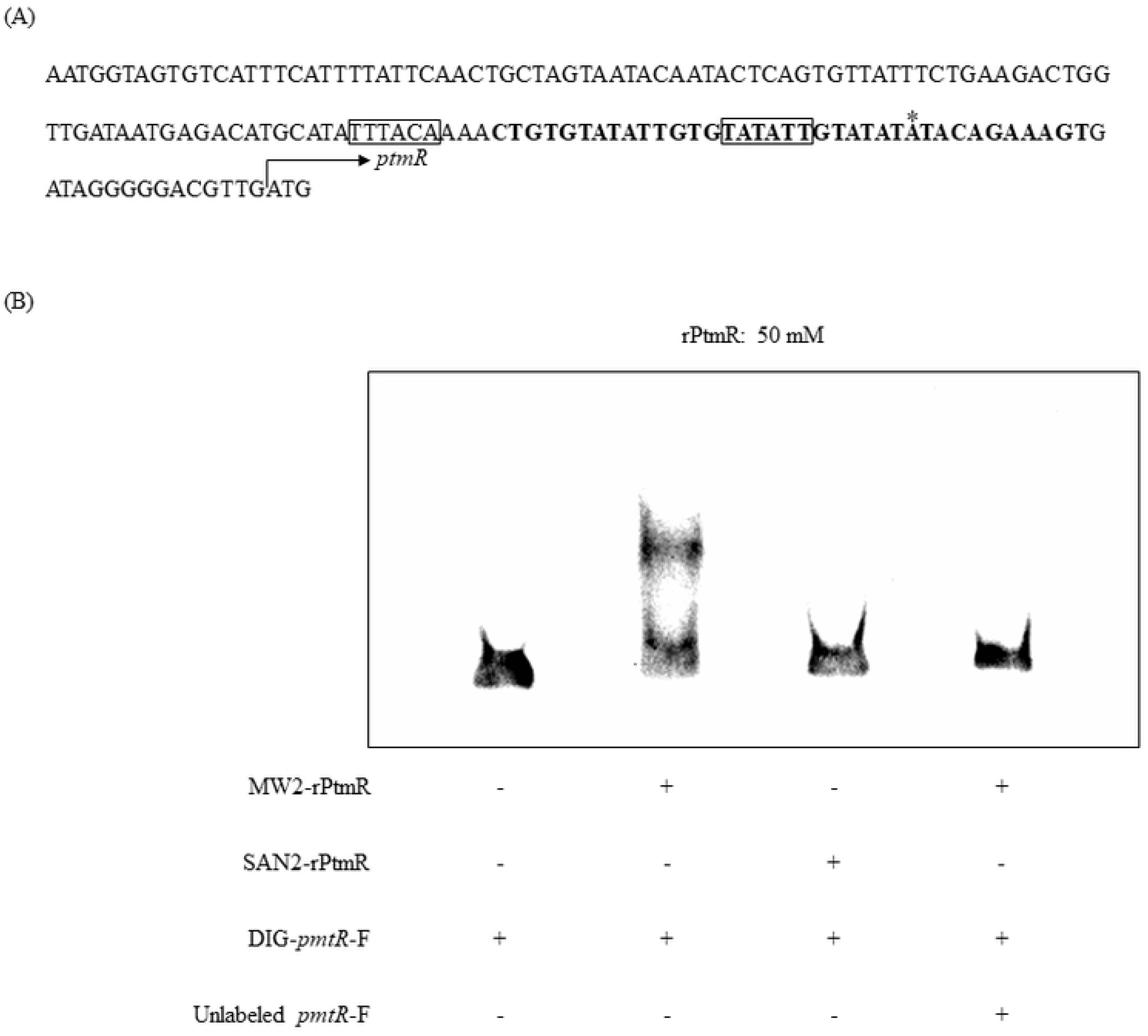
EMSAs of PmtR. (A) The nucleotide sequence of the *pmtR* promoter region. Squares, −35, −10 box; *, *pmtR* transcriptional start site; bold, PmtR binding region. (B) EMSAs of PmtR with the DNA fragment labeled with DIG were performed with the method described in the Materials and Methods.

### Hemolytic activity in the MW2, SAN2 and MM2278

Since PSM transported by PmtA-D has been demonstrated to be involved in hemolytic activity (32), we thought that overexpression of PmtA-D affected the hemolytic activity, we next analyzed hemolytic activities of strains MW2 and SAN2 on sheep blood agar. Compared to MW2 wild type, SAN2 produced a larger hemolytic zone (Fig. 5A). When *pmtR* and *pmtA-D* (*pmtR-D*: MW1875 to MW1871) were inactivated in SAN2 (MM2278), the hemolytic zone became smaller than that of SAN2. The hemolytic zone produced by the complement stain MM2279 was similar in size as that of SAN2 (Fig. 5A).

**Fig. 5.**
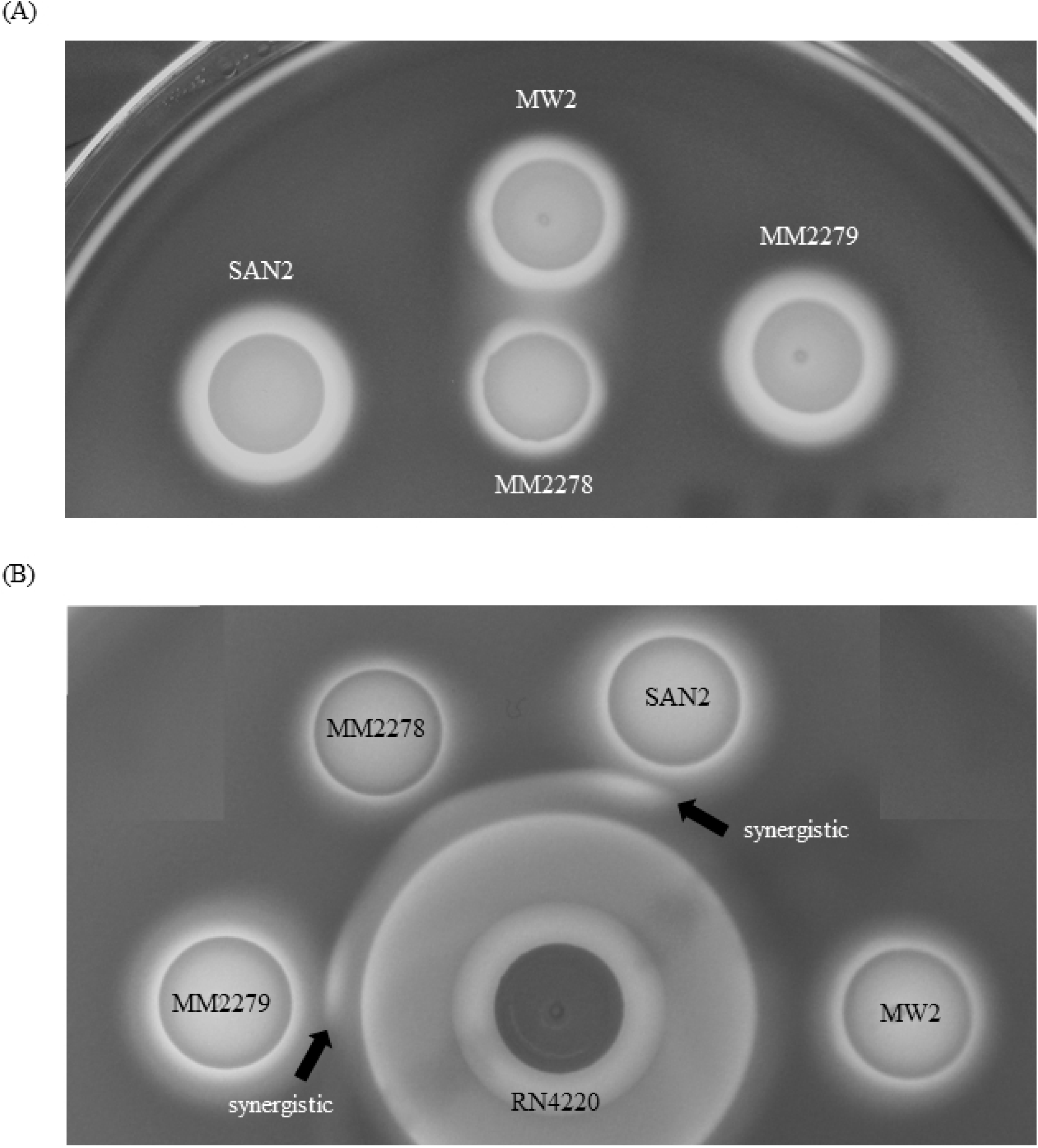
Hemolytic assay of *S. aureus* mutants. Ten microliters of a 10-fold dilution of an *S. aureus* overnight culture was spotted on a sheep blood agar plate. Agar plates were incubated for 20 h and then kept at 4°C for 2 days.

It was reported that PSM or delta-hemolysin enhanced the activity of beta-hemolysin (33). Since MW2 (and SAN2) do not produce beta-hemolysin, we investigated a synergy hemolytic activity of strains SAN2 and RN4220 (which produces beta-hemolysin). An enhanced hemolytic zone was produced (black arrows) by SAN2 and RN4220, while MW2 and MM2278 did not have an enhanced hemolytic effect with RN4220 (Fig. 5B).

### Susceptibility to LL37 and hBD3 in theMW2, SAN2, MM2278 and MM2279

Since it was reported that *pmtR-D* inactivation increased the susceptibility to LL37 and hBD3 in a previous study (34), we investigated the effect of overexpression of PmtA-D in SAN2 on the susceptibility to hBD3 (Fig. 6) and LL37 (supplemental Fig. 3). As shown in Fig. 6, compared to the MW2 original strain, SAN2 showed a decrease in susceptibility to hBD3, while the inactivation of *pmtR-D* increased susceptibility to the peptides, resulting in the same susceptibility as the MW2. The complement strain showed similar susceptibility with that of SAN2.

**Fig. 6.**
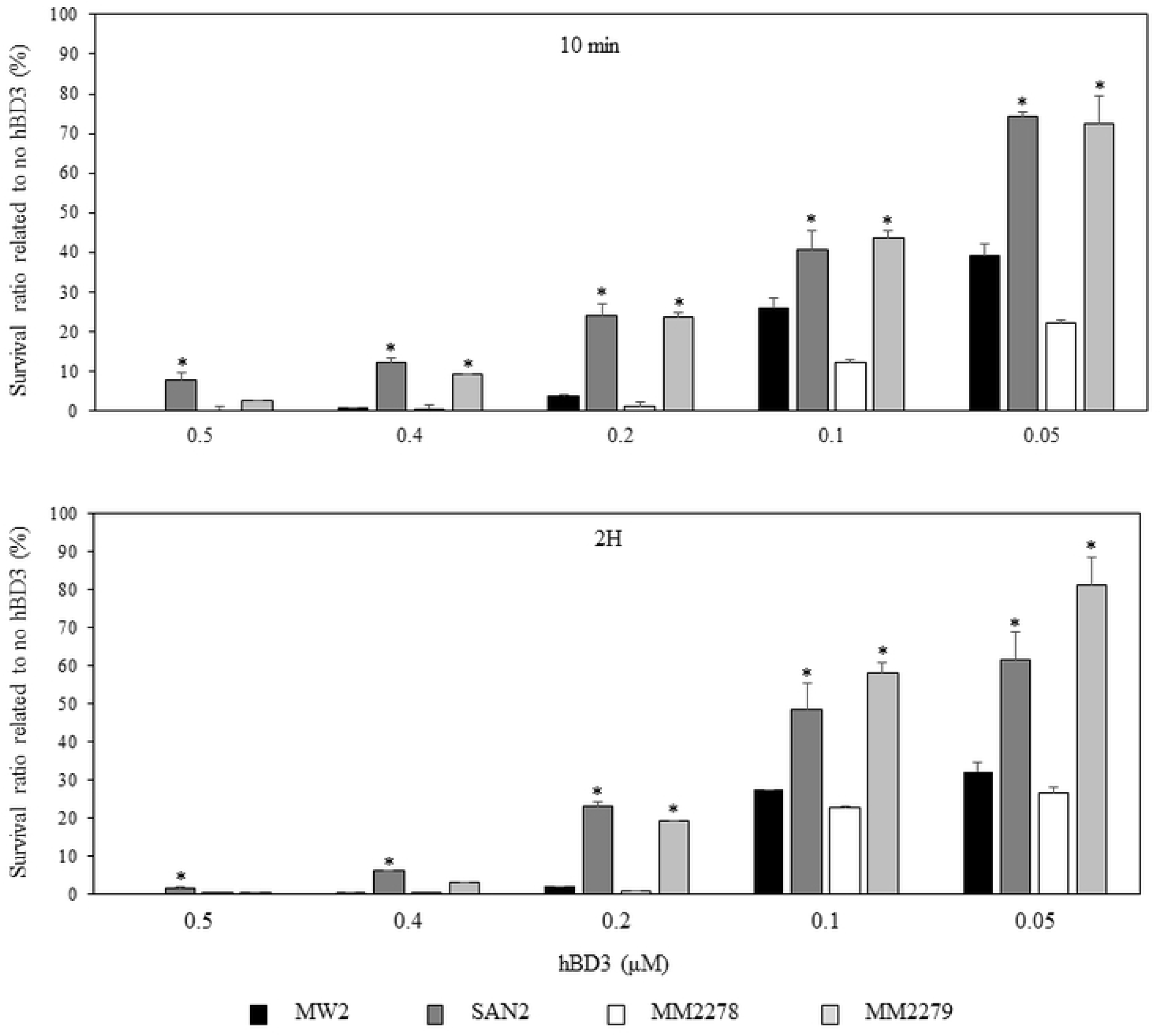
Susceptibility of *S. aureus* mutants to human defensin 3. (A) An antibacterial assay for hBD3 was performed as described elsewhere. Various concentrations of hBD3 (0.05 to 0.5 μM) were applied to a bacterial suspension (10^5^ cells per 500 μl phosphate buffer). After 10min or 2 h, dilutions of the reaction mixture were plated on agar media. After incubation at 37°C overnight, the colony-forming units (CFUs) were counted. The antibacterial effect was calculated as the ratio of the number of surviving cells (survival rate %) to the total number of bacteria incubated in control PB solution after exposure to antimicrobial peptides. (B) An antibacterial assay with 0.6 μM LL37 was performed. \**p* <0.05 as determined increase by Dunnett’s post hoc tests compared to untreated MW2.

### Mice survival rate with injection of the MW2, SAN2 and MM2278

Finally, we performed the mouse survival experiment using the bacteremia model. Injection of the MW2 strain killed only one mouse out of the 8 mice 3 days after injection, while 5 of 8 mice were killed between 1 to 5 days after the injection of SAN2 (Fig. 7A, P=0.010, log-rank test). No mouse was killed by the injection of the SAN2 *pmtR-D* inactivation strains (MM2278) (Fig. 7B, P=0.002, log-rank test).

**Fig. 7.**
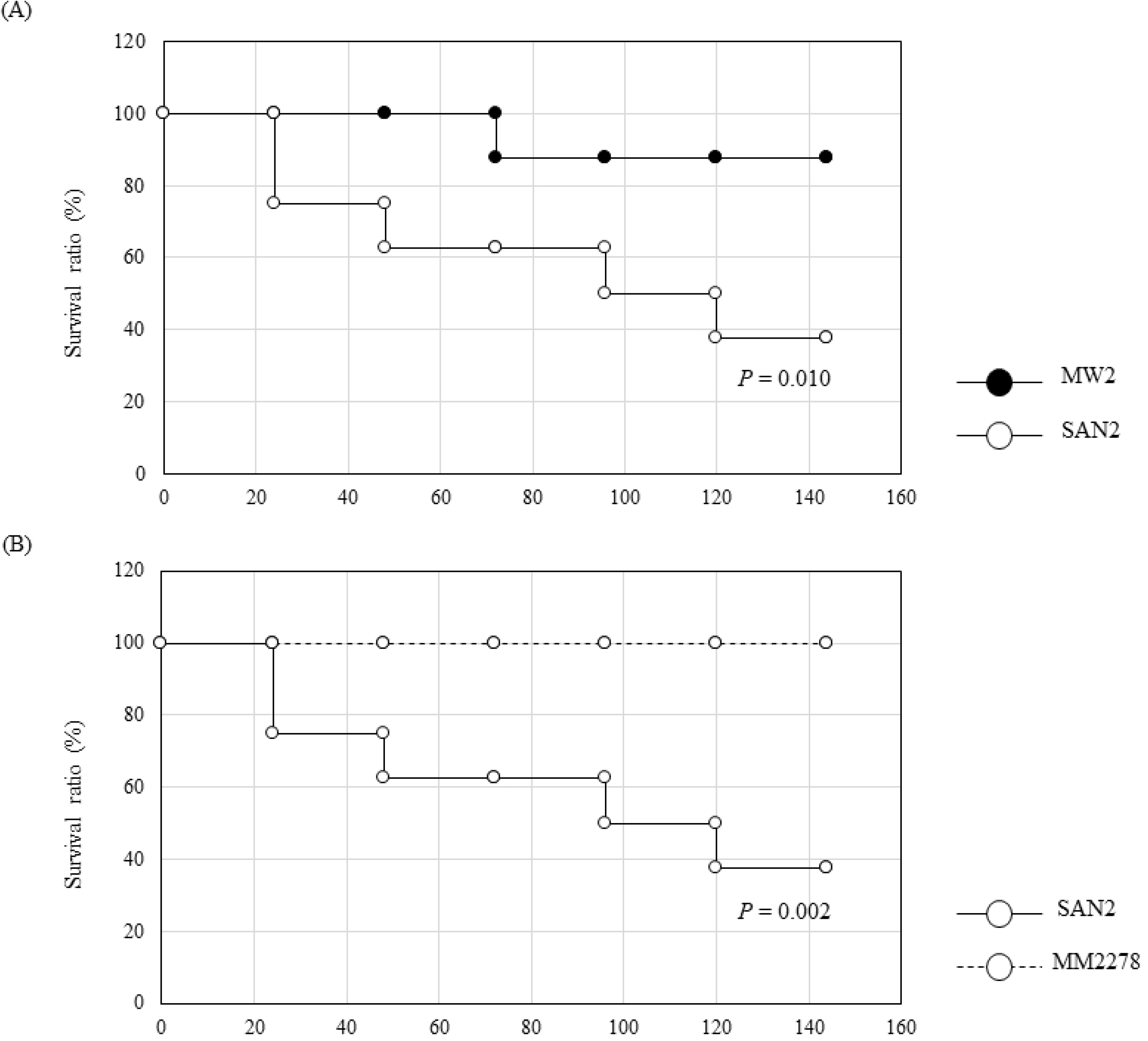
Mouse survival experiment. Survival percentage of Slc:ddY mice after being challenged with intravenous injection of 1.5 × 10^8^ CFU of *S. aureus*. (A) Survival comparison between MW2 and SAN2. Number of trials: MW2=8, SAN2=8. Survival statistics were calculated by log-rank test. Significant differences compared with MW2 are shown. (B) Survival comparison between SAN2 wild type and pmtR,A-D inactivation mutant of SAN2 (MM2278). Number of trials: SAN2=8, MM2278=6. Survival statistics were calculated by log-rank test. Significant differences compared with SAN2 are shown.

## Discussion

In this study, we first demonstrated a new highly nisin A resistant mechanism independent from BraRS-VraDE system. We obtained 4 PmtA-D-overexpressing mutant strains from MW2, COL and TY34 that acquired high nisin A resistance. All these mutants had a point mutation in the *pmtR* gene, yielding a mutant PmtR with an Ala43D substitution (SAN2 from MW2) or truncated PmtR (SAN469 from MW2, SAN233 from COL and SAN455 from TY34). EMSAs showed that the SAN2-derived PmtR protein (SAN2-rPmtR) did not bind the DNA region upstream of *pmtR-D*. Since PmtR is a negative transcriptional regulator of the *pmtRABCD* operon (31), SAN2-PmtR and the other three truncated PmtRs could not suppress the expression of the *pmtR-D* operon, resulting in the overexpression of *pmtR-D*. The mutation site of PmtR in SAN2 is within helix-turn-helix DNA binding region. Based on our EMSA assay, a mutated-PmtR in SAN2 lost the binding ability to the target DNA region, suggesting that the mutation site is the critical for DNA binding or mutated-PmtR in SAN2 occuers a structural change. PmtA-D form an ABC transporter from two membrane proteins (PmtA and C) and two ATPases (PmtB and D) (30). PmtA-D is involved in the transportation of PSMs and delta-hemolysin (Hld) from the cytoplasm to the extracellular space (30, 32). PSMs have versatile virulence activity such as surface spreading activity responsible for epithelial colonization, biofilm formation, proinflammatory activity, cytolytic activity and antimicrobial activity (35, 36). In addition, Cheung GYC et al. have recently reported that the Pmt transporter is also associated with human-derived antimicrobial peptides such as hBD3 and LL37 (34). We also found that high expression of Pmt transporters in SAN2 resulted in high resistance against LL37 and hBD3. Although there are no clear structural similarities among PSMs, delta-hemolysin, hBD3 and LL37, all these peptides have membrane insertional activity. Based on these findings, we consider nisin A, which also has membrane insertional activity, to be exported by the Pmt system. Therefore, the high nisin A resistance in SAN2, SAN469, SAN233 and SAN455 is mediated by overexpression of PmtA-D. However, we also evaluated the susceptibility to gallidermin, bacitracin and nukacin ISK-1 and found no difference between MW2 and SAN2 strains. Nukacin ISK-1 is functionally close to nisin A because these are lantibiotics with the same target, lipid II (4, 8). These results indicate that the Pmt system recognizes the limited antimicrobial peptides with membrane insertional activity.

Joo HS et al. demonstrated that PSMα1-3 binds PmtR to release it from its target DNA region, which is followed by induction of the expression of the *pmt* operon (31). PSM expression is upregulated by the *agr* system (32, 35). Since *agr* expression increases from the late exponential phase (37, 38), the expression levels of PSM and PmtA-D are increased at the late exponential phase. In contrast, in the SAN2 mutant, PmtA-D expression was independent of the growth phase due to the lack of functional PmtR (data not shown). Therefore, PmtA-D are constitutively overproduced, thus PSMs are constantly exported to outside the cell during SAN2 growth. As mentioned above, PSMs have versatile virulent activities, so the increased PSM transport may modulate the virulence of *S. aureus*. To support this, SAN2 showed highly pathogenic. We found that the hemolytic activity of SAN2 was greater than that of the MW2 wild type because PSMα has an enhanced effect on hemolytic activity (33). Additionally, the susceptibility to human antimicrobial peptides was decreased in the mutants. Finally, we found in a mouse experiment that compared to the MW2 wild type, SAN2 showed an increase in fatality rate. The Pmt-dependent nisin A resistance identified in this study is different from the previously identified BraRS-VraDE system in terms of the modulation for virulence in *S. aureus*.

Previously, we isolated strains highly resistant to nisin A that showed highly constitutive expression of VraDE (25). VraDE, which is regulated by BraRS, is an intrinsic resistance factor against nisin A, gallidermin, nukacin ISK-1 and bacitracin. In MW2, sub-MIC (1/16 MIC) nisin A induced the expression of VraDE, while sub-MIC nisin A did not induce its expression in SAN2. However, *pmtA-D* inactivation in SAN2 caused inducible expression of VraDE by sub-MIC nisin A. The induction of *vraD* expression occurred at high concentrations of nisin A in SAN2, suggesting that the Pmt system is an dominant system contribute to nisin A resistance in SAN2.

In conclusion, we found a new mechanism for high nisin A resistance that is mediated by the overexpression of Pmts in *S. aureus*. It is important to note that Pmt overexpression-dependent high nisin A resistance also enhances the virulence. Since nisin A is widely used as a food additive, it is important to be careful in its use.

## Materials and Methods

### Ethics statement

The animal experimentation was conducted according to the protocol approved by the President of Kagoshima University after the review by the Institutional Animal Care and Use Committee (Ethical number: D18015).

### Bacterial strains and growth conditions

The bacterial strains used in this study are shown in Table 1. *S*. *aureus* and *Escherichia coli* XL-II were grown in Trypticase soy broth (TSB) (Beckton Dickinson Microbiology Systems, Cockeysville, MD, USA) and Luria-Bertani (LB) broth, respectively. Tetracycline (TC, 5 µg/ml) and chloramphenicol (5 µg/ml) were used for *S. aureus*, and ampicillin (100 µg/ml) was used for *E. coli* when necessary.

**Table1.**
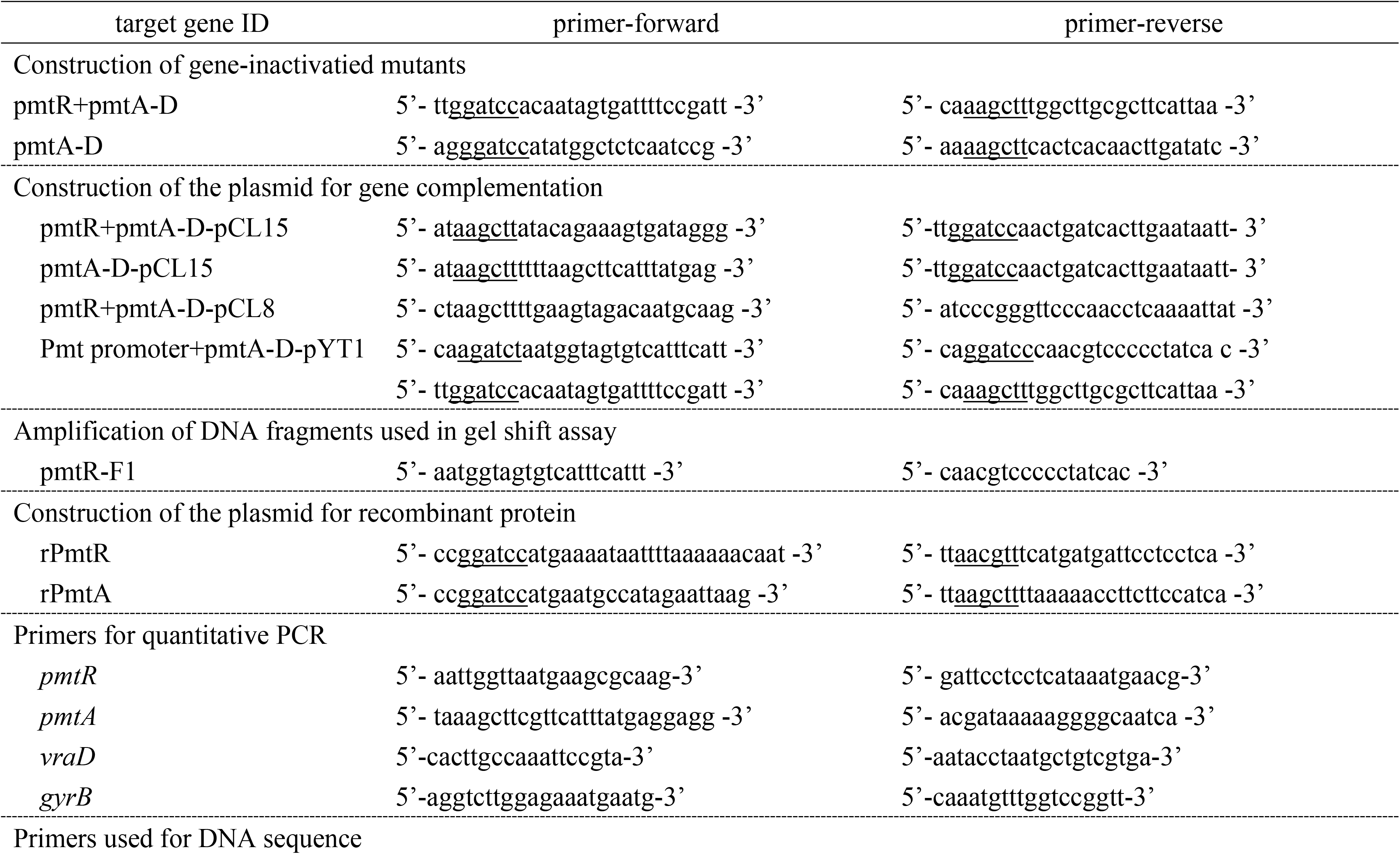
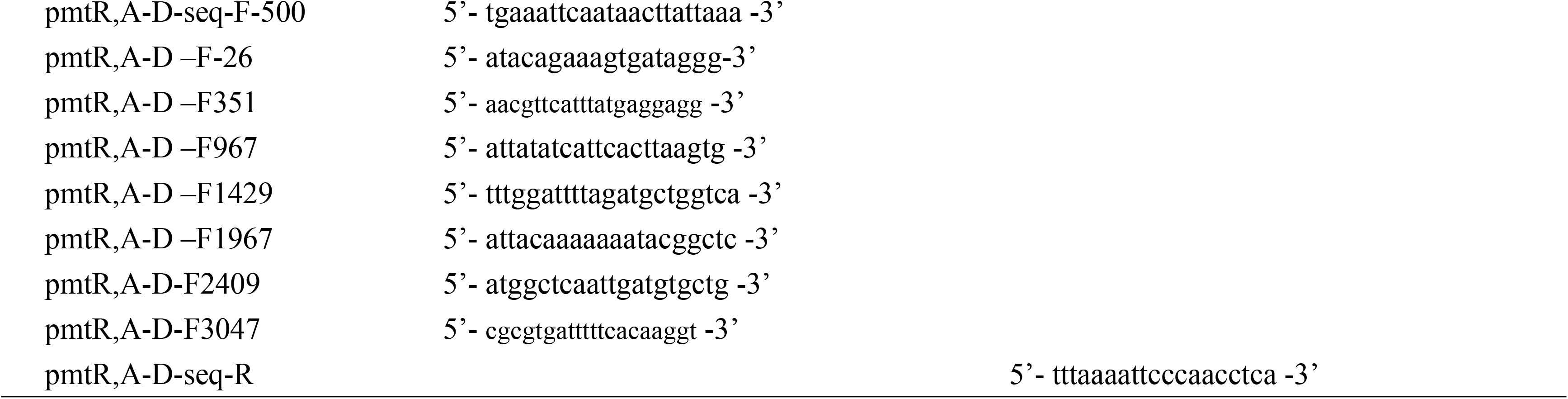
primers used in this study.

### Isolation of mutants highly resistant to nisin A

*S. aureus* mutants highly resistant to nisin A were obtained by a method described elsewhere (25). Briefly, a microdilution method that is generally used to evaluate the minimum inhibitory concentration (MIC) of antibacterial agents was used for isolation of the mutants. Overnight culture of *S. aureus* MW2 was diluted to 10^7^ cells/ml, and 10 µl of the dilute culture was applied to each well (100 µl), which contained a 2-fold dilution series of nisin A (16 to 16384 µg/ml). After incubation at 37°C overnight, bacterial cells that grew at 1/2 MIC nisin A were taken and diluted 100 fold. Ten microliters of this diluted sample was then applied to each well (100 µl), which contained serial 2-fold dilutions of nisin A (16 to 16384 µg/ml). This procedure was repeated three times. At the end, bacterial cells grown in the presence of 1/2 MIC nisin A were appropriately diluted and plated on tryptic soy agar (TSA). After overnight incubation, 14 colonies were picked up and incubated in 5 ml of TSB. Overnight culture was used for the determination of the MIC of nisin A. This experiment was independently performed 3 times. Furthermore, we tried to isolate highly nisin A-resistant mutants from other *S. aureus* strains.

We also tried to isolate strains highly resistant to nisin A from *S. aureus* COL and TY34 by the same method described above.

### Microarray analysis

Overnight cultures of *S. aureus* (10^8^ cells) were inoculated into 30 ml of fresh TSB and cultured at 37°C with shaking. When the OD_660_ reached 0.4, the bacterial cells were collected by centrifugation at 5,000 x *g* for 5 min at 4°C. Total RNA was extracted by using the FastRNA Pro Blue Kit (MP Biomedicals, Cleveland, OH, USA) according to the manufacturer’s protocol. Then, cDNA of each sample was synthesized from 10 μg of total RNA using the FairPlay III Microarray Labeling Kit (Agilent Technologies, Santa Clara, CA, USA) according to the manufacturer’s instructions. The Agilent eArray platform (Agilent Technologies) was used to design the microarray: 13,939 probes (60-mers) were designed to target the 2,628 protein-coding genes of *S. aureus* MW2 (up to five probes per gene). Microarray analysis was performed by a method described elsewhere. The experiments were performed using two biological replicates (two technical replicates for each set of conditions), and the expression data were deposited into the Gene Expression Omnibus (http://www.ncbi.nlm.nih.gov/geo/) under accession GSE131352.

### DNA sequences of MW1875-1871 regions

Since we found the increased expression of genes corresponding to MW1875-1871 in the SAN2 strain, we determined the DNA sequence of the MW1875-1871 regions. Primers were constructed to amplify MW1875-1871 with their corresponding flanking regions, including the promoter regions of MW1875 (Table 1). To prepare chromosomal DNA from the MW2 original strain and the mutant strains, the cells from 1 ml of overnight cultures were collected. The cells were suspended in 100 µl of 10 mM Tris-HCl (pH 6.8) containing 10 µg of lysostaphin (Sigma Aldrich), incubated at 37°C for 20 min, and then incubated at 95°C for 15 min. After centrifugation, cell lysates were used as template DNA for PCR. PCR was performed using the Takara Ex Taq system, and the amplicons were purified using a QIAquick kit (QIAGEN, Hilden, Germany). The nucleotide sequences of each DNA fragment were determined using specific primers. The primers used to amplify DNA sequences are listed in Table 1.

### Inactivation of MW1875 and MW1874 in *S. aureus* and their complementation

Strains used in this study were listed in table 2. The specific gene inactivation by the insertion of thermosensitve plasmid pYT1 was performed by a method described elsewhere (26). An internal DNA fragment of MW1875 or MW1874 was amplified by PCR using specific primers, and then cloned into a pYT1 vector. The plasmid was electroporated into *S. aureus* RN4220, and then the plasmid in RN4220 was transferred into respective strains by transduction using phage 80α. The obtained strains were grown overnight at 30°C. The appropriate dilutions of the culture were poured on TSA plates containing TC (10 µg/ml) and then incubated at 42°C overnight. Colonies were picked up and replated on TSA containing TC. The disruption of the target gene was checked by PCR. For gene complementation, the vector pCL15, which has an isopropyl-β-d-thiogalactopyranoside (IPTG)-inducible promoter upstream of the cloning site, was used. A DNA fragment for complementation was PCR-amplified using chromosomal DNA from the MW2 or mutant strains. The DNA fragment was cloned into pCL15 using *E. coli* XL-II competent cells. The obtained plasmid was electroporated into *S. aureus* RN4220 and was subsequently transduced into the appropriate strain using the phage 80α.

**Table 2.**
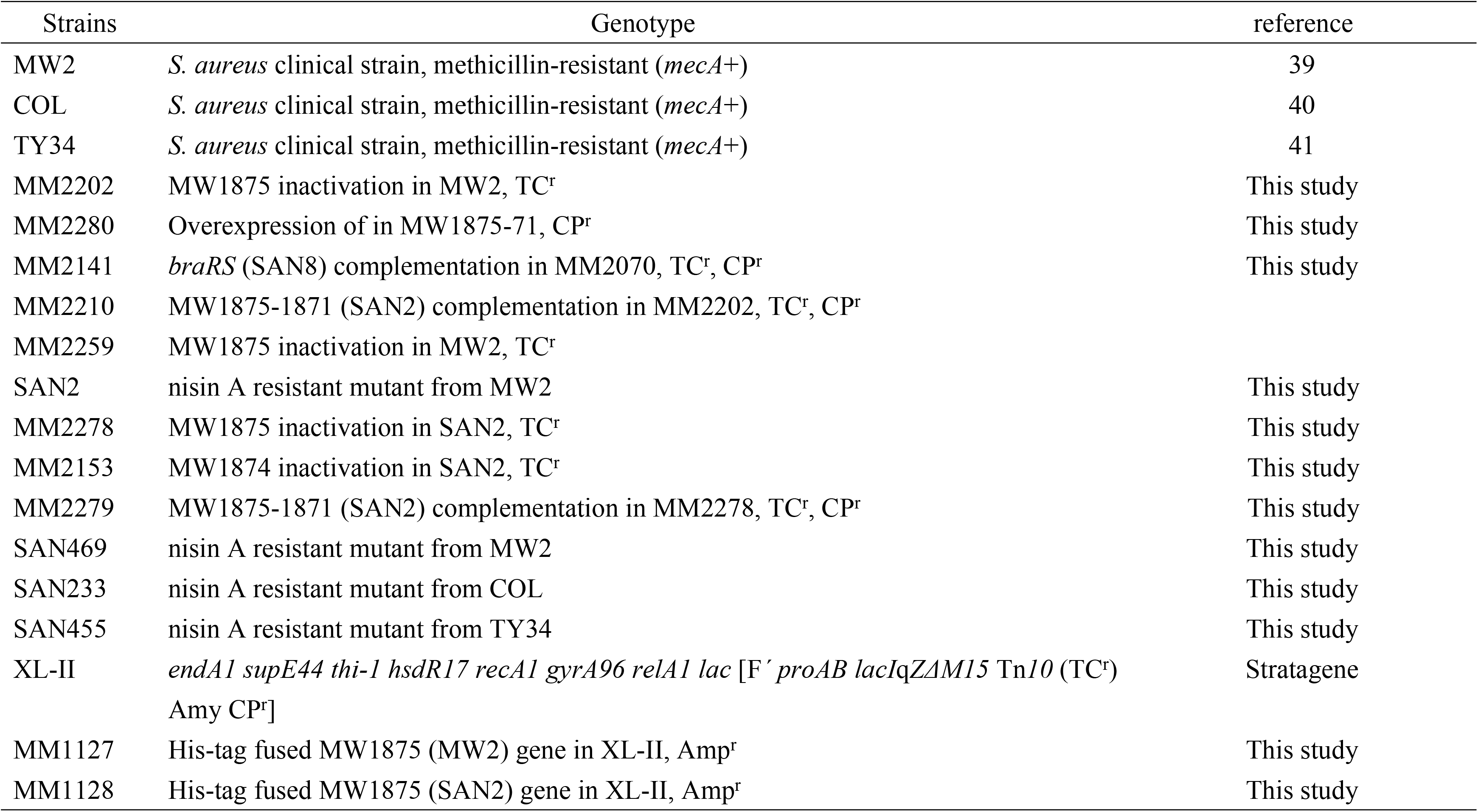
strains used in this study.

**Table3.** susceptibility of nisin A, galidermin, nukacin ISK-1, and bacitracin against MW2, COL, TY34 and individual nisin A-resistant SAN strains.

|  | nisin A | BC | galidermin | nukacin<br>ISK-1 |
| --- | --- | --- | --- | --- |
| MW2 | 512 | 64 | 8 | 16 |
| SAN2 | 2048 | 64 | 8 | 16 |
| SAN469 | 2048 | 64 | 8 | 16 |
| COL | 1024 | 64 | 8 | 16 |
| SAN233 | 2048 | 64 | 8 | 16 |
| TY34 | 512 | 64 | 8 | 16 |
| SAN455 | 2048 | 64 | 8 | 16 |

BC, bacitracin

To obtain a mutant that constitutively expresses MW1874-1871 by their own promoter upstream of MW1875 without MW1875 expression, we constructed a plasmid by PCR cloning a DNA fragment containing the internal region of MW1875 (295 bp) and the promoter region of MW1875-1871 (149 bp). Two PCR fragments were obtained and then cloned into the pYT1 vector by using restriction enzymes. The plasmid was finally transduced into the MW2 strain. The plasmid integration was performed with the method described above. Two types of the mutants were obtained (Suppl. Fig. 1).

Finally, the strain that contained inactivated MW1875 and expressed MW1874-1871 was verified by PCR and DNA sequencing.

### Quantitative PCR and immunoblotting analysis

Quantitative PCR was performed to investigate the expression of MW1875 (*pmtR*), MW1874 (*pmtA*) and *vraD*. A small portion of overnight culture (10^8^ cells) was inoculated into 5 ml of fresh TSB and then grown at 37°C with shaking. When the optical density at 660 nm reached 0.5, nisin A (32 µg/ml) or bacitracin (32 µg/ml) was added to the bacterial culture. After incubating for 15 min (for quantitative PCR) and 2 h (for immunoblotting), the bacterial cells were collected. For quantitative PCR, RNA extraction, cDNA synthesis and PCR were performed as described previously (25).

For immunoblotting, antiserum against MW1875 and MW1874 was obtained by immunizing rabbit with the recombinant protein as described previously (27). Briefly, the coding region of MW1875 or MW1874 amplified by PCR using specific primers was cloned into pQE30 (QIAGEN, Tokyo, Japan) which is used for the construction of histidine-tagged recombinant proteins. The obtained plasmid was transformed into *E. coli* M15 (pREP4). The resulting recombinant protein was purified according to the manufacturer’s instructions. Antiserum was obtained using recombinant protein. We also immunoblotted VraD using anti-rVraD obtained previously.

The collected bacterial cells were resuspended in 200 µl of Tris-HCl (pH 6.8) containing 10 µg of lysostaphin and incubated for 20 min at 37°C. The cells were then heated at 95°C for 15 min. After centrifugation, the supernatant was obtained as a whole-cell lysate. Lysate proteins mixed with an equal volume of sample loading buffer were resolved by 15% SDS-polyacrylamide gel electrophoresis (PAGE). Then, the proteins were transferred to a nitrocellulose membrane. After blocking with 2% skim milk in Tris-buffered saline (TBS; 20 mM Tris, 137 mM NaCl [pH 8.0]) containing 0.05% Tween 20 (TBS-T), the membrane was incubated with specific antiserum (diluted 1:1,000 in 1% skim milk in TBS-T) for 1 h at 37°C. The membrane was then washed with TBS-T and incubated with horseradish peroxidase-conjugated anti-mouse IgG (diluted 1:1,000 in TBS-T) (Promega, Madison, WI, USA) for 1 h at 37°C. The membrane was then washed 5 times with TBS-T, and the protein band reacting with the antiserum was detected using a chemiluminescence detection system (PerkinElmer, Waltham, MA, USA).

### Hemolysis assay

Hemolytic assays were performed by using TSB-containing 5% sheep blood agar plates (Becton, Dickinson and Company, Japan). Three microliters of overnight culture was spotted on the agar plate. Agar plates were incubated for 20 h and then kept at 4°C for 2 days.

### Susceptibility test of LL37 and human defensin 3

An antibacterial assay was performed as described elsewhere (28). Briefly, an overnight culture of *S. aureus* strains was collected and washed with 10 mM sodium phosphate buffer (PB). The bacterial suspension was diluted to 10^7^ cells/ml with PB, and 10 µl of the bacterial suspension (10^5^ cells) was inoculated into 500 µl of PB with or without human antimicrobial peptides (beta-defensin-3; Peptide Institute Inc., Osaka, Japan, or LL37; 0.6 μM) and incubated aerobically for 2 h at 37°C. Dilutions of the reaction mixture (100 µl) were plated on agar media and incubated at 37°C overnight. The colony-forming units (CFUs) were determined as the total number of colonies on each plate. The antibacterial effect was calculated as the ratio of the number of surviving cells (survival rate %) to the total number of bacteria incubated in control PB solution after exposure to antimicrobial peptides.

### Electrophoretic mobility shift assay (EMSA)

For the EMSA, two 6× histidine-tagged recombinant PmtR (rPmtR) proteins from the MW2 original strain and SAN2 were used.For construction rPmtRs were performed as described above. Briefly, a DNA fragment encoding *pmtR* (MW1875) was amplified with specific primers by using chromosomal DNA of the MW2 and SAN2. Then, DNA fragments were subsequently cloned into pQE30 (QIAGEN, Tokyo, Japan). The plasmid was then transformed into *E. coli* M15 (pREP4). The recombinant protein was purified according to the manufacturer’s instructions. To assess the binding of rPmtR to a region upstream of *pmtR*, an EMSA was performed as described previously (25). A DNA fragment encompassing the region upstream of *pmtR, A-D* was amplified with the specific primers listed in Table 1. The DNA fragments were labeled at the 3′ end with digoxigenin (DIG) using a DIG Gel Shift Kit, 2^nd^ Generation (Roche, Mannheim, Germany). The DIG-labeled fragment (5 ng) was reacted with rPmtR protein (50 mM) in the labeling buffer provided with the kit. When necessary, a nonlabeled DNA fragment (10 ng) was added to the reaction mixture. After native PAGE on a 6% polyacrylamide gel, the DNA fragments were transferred to a positively charged nylon membrane (Roche, Mannheim, Germany) and visualized according to the manufacturer’s protocol.

### Mouse bacteremia experiment

A mouse bacteremia experiment was performed as described previously (29). Six-week-old female Slc:ddY mice were purchased from SLC (Shizuoka, Japan). Small portions of overnight cultures of *S. aureus* MW2 original strain, SAN2 and SAN2 with MW1875-1871 inactivation (MM2278) were inoculated into 5 ml of fresh TSB and incubated at 37°C with shaking. When the OD_660_ reached 0.8, bacterial cells were collected and washed with PBS. Then, the cells were resuspended in PBS at a concentration of 1.0 × 10^9^ CFU/ml. An aliquot of 100 μl (1.0 × 10^8^ CFU) was injected into the tail vein of the mice. Mouse survival was monitored for 6 days. This experimental was approved by the ethics committee of the Animal Research Center of Kagoshima University, Japan.

## Acknowledments

This study was supported in part by Grant-in-Aid for Scientific Research (C) (Grant No: 18K09553) from the Ministry of Education, Culture, Sports, Sciences, and Technology of Japan.

## Supporting information legends

**Fgure S1. Construction of the strain which had a MW1875-inactivation and expressed MW1874-1871.**

To obtain a mutant that constitutively expresses MW1874-1871 by their own promoter upstream of MW1875 without MW1875 expression, we constructed a plasmid containing the internal region of MW1875 (295 bp) and the promoter region of MW1875-1871 (149 bp). The plasmid was finally transduced into the MW2 strain. The plasmid integration was performed with the method described Materials and Methods section. Two types of the mutants (Case 1and 2) were obtained and the mutant with Case1 was selected for further experiments.

**Fgure S2. Expressions of MW1875 and MW1874 by addition of nisin A in MW2 and SAN2**

Expression of *pmtR and pmtA* in *S. aureus* MW2 and SAN2 by addition of various concentrations of nisin A was evaluated by quantitative PCR. Exponential phase cells that reached OD_660_=0.5 were exposed to various concentrations of nisin A. After incubation for 15 min, the bacterial cells were collected. RNA extraction, cDNA synthesis and quantitative PCR were performed as described in Materials and Methods section. Statistical analysis was determined by Dunnett’s post hoc tests compared to untreated MW2 or SAN2.

**Fgure S3. Susceptibility of *S. aureus* mutants to LL37**

An antibacterial assay for LL37 was performed as described in Materials and Methods section. LL37 (0.6 μM) were applied to a bacterial suspension (10^5^ cells per 500 μl phosphate buffer). After 10min or 2 h, dilutions of the reaction mixture were plated on agar media. After incubation at 37°C overnight, the colony-forming units (CFUs) were counted. The antibacterial effect was calculated as the ratio of the number of surviving cells (survival rate %) to the total number of bacteria incubated in control PB solution after exposure to antimicrobial peptides. \**p* <0.05 as determined increase by Dunnett’s post hoc tests compared to untreated MW2.

## References

1. Cotter PD, Hill C, Ross RP. Bacteriocins: developing innate immunity for food. Nat Rev Microbiol. 2005; 3: 777–788.

2. Jack RW, Tagg JR, Ray B. Bacteriocins of gram-positive bacteria. Microbiol. Rev. 1995; 59: 171–200.

3. Nissen-Meyer J, Nes IF. Ribosomally synthesized antimicrobial peptides: their function, structure, biogenesis, and mechanism of action. Arch Microbiol 1997; 167: 67–77.

4. Nishie M, Nagao J, Sonomoto K. Antibacterial peptides “bacteriocins”: an overview of their diverse characteristics and applications. Biocontrol Sci. 2012; 17(1): 1–16.

5. Nagao J, Asaduzzaman SM, Aso Y, Okuda K, Nakayama J, Sonomoto K. Lantibiotics: insight and foresight for new paradigm. J Biosci Bioeng. 2006; 102: 139–149.

6. Nes IF, Holo H. Class II antimicrobial peptides from lactic acid bacteria. Biopolymers 2000; 55: 50–61.

7. Delves-Broughton J, Blackburn P, Evans RJ. and Hugenholtz, J. Applications of the bacteriocin, nisin. Antonie Van Leeuwenhoek. 1996; 69: 193–202.

8. Bierbaum G, Sahl HG. Lantibiotics: mode of action, biosynthesis and bioengineering. Curr Pharm Biotechnol. 2009; 10: 2–18.

9. Lubelski J, Rink R, Khusainov R, Moll GN, Kuipers OP. Biosynthesis, immunity, regulation, mode of action and engineering of the model lantibiotic nisin. Cell Mol Life Sci. 2008; 65: 455–476.

10. Shin JM, Gwak JW, Kamarajan P, Fenno JC, Rickard AH, Kapila YL. Biomedical applications of nisin. J Appl Microbiol. 2016; 120(6): 1449–65.

11. Le Lay C, Dridi L, Bergeron MG, Ouellette M, Fliss IL. Nisin is an effective inhibitor of *Clostridium difficile* vegetative cells and spore germination. J Med Microbiol. 2016; 65(2): 169–75.

12. Udompijitkul P, Paredes-Sabja D, Sarker MR. Inhibitory effects of nisin against Clostridium perfringens food poisoning and nonfood-borne isolates. J Food Sci. 2012; 77(1): M51–6.

13. Tong Z, Ni L, Ling J. Antibacterial peptide nisin: a potential role in the inhibition of oral pathogenic bacteria. Peptides. 2014; 60: 32–40.

14. Okuda K, Zendo T, Sugimoto S, Iwase T, Tajima A, Yamada S, et. al. Effects of bacteriocins on methicillin-resistant *Staphylococcus aureus* biofilm. Antimicrob Agents Chemother. 2013; 57(11): 5572–9.

15. Field D, O’ Connor R, Cotter PD, Ross RP, Hill C. In Vitro Activities of Nisin and Nisin Derivatives Alone and In Combination with Antibiotics against Staphylococcus Biofilms. Front Microbiol. 2016; 18: 7:508.

16. Gharsallaoui A, Oulahal N, Joly C, Degraeve P. Nisin as a Food Preservative: Part 1: Physicochemical Properties, Antimicrobial Activity, and Main Uses. Crit Rev Food Sci Nutr. 2016; 56: 1262–74.

17. Juturu V, Wu JC. Microbial production of bacteriocins: Latest research development and applications. Biotechnol Adv. 2018; 36(8): 2187–2200.

18. van Harten RM, Willems RJL, Martin NI, Hendrickx APA. Multidrug-Resistant Enterococcal Infections: New Compounds, Novel Antimicrobial Therapies?Trends Microbiol. 2017; 25(6): 467–479.

19. Lowy FD. *Staphylococcus aureus* infections. New Engl J Med. 1998; 339: 520–532.

20. Manders SM. Toxin-mediated streptococcal and staphylococcal disease. J Am Acad Dermatol. 1998; 39: 383–388.

21. Foster TJ. The *Staphylococcus aureus* “superbug”. J Clin Invest. 2004; 114: 1693–1696.

22. Silva CCG, Silva SPM, Ribeiro SC. Application of Bacteriocins and Protective Cultures in Dairy Food Preservation. Front Microbiol. 2018; 9: 9:594.

23. Kawada-Matsuo M., Yoshida Y, Zendo T, Nagao J, Oogai Y, Nakamura Y, et al. Three distinct two-component systems are involved in resistance to the class I bacteriocins, Nukacin ISK-1 and nisin A, in *Staphylococcus aureus*. PLoS One. 2013; 8: e69455.

24. Hiron A, Falord M, Valle J, Débarbouillé M, Msadek T. Bacitracin and nisin resistance in *Staphylococcus aureus*: a novel pathway involving the BraS/BraR two-component system (SA2417/SA2418) and both the BraD/BraE and VraD/VraE ABC transporters. Mol Microbiol. 2011; 81: 602–22.

25. Arii K, Kawada-Matsuo M, Oogai Y, Noguchi K, Komatsuzawa H. Single mutations in BraRS confer high resistance against nisin A in *Staphylococcus aureus*. Microbiologyopen. 2019; 17: e791.

26. Kajimura J, Fujiwara T, Yamada S, Suzawa Y, Nishida T, Oyamada Y, et. al. Identification and molecular characterization of an *N*-acetylmuramyl-l-alanine amidase Sle1 involved in cell separation of *Staphylococcus aureus*. Mol Microbiol. 2005; 58: 1087–1101.

27. Kawada-Matsuo M, Oogai Y, Zendo T, Nagao J, Shibata Y, Yamashita Y, et al. Involvement of the novel two-component NsrRS and LcrRS systems in distinct resistance pathways against nisin A and nukacin ISK-1 in *Streptococcus mutans*. Appl Environ Microbiol. 2013; 79: 4751–4755.

28. Matsuo M, Oogai Y, Kato F, Sugai M, Komatsuzawa H. Growth-phase dependence of susceptibility to antimicrobial peptides in *Staphylococcus aureus*. Microbiology. 2011; 157(Pt 6): 1786–97.

29. Oogai Y, Yamaguchi M, Kawada-Matsuo M, Sumitomo T, Kawabata S, Komatsuzawa H. Lysine and Threonine Biosynthesis from Aspartate Contributes to *Staphylococcus aureus* Growth in Calf Serum. Appl Environ Microbiol. 2016; 82(20): 6150–6157.

30. Chatterjee SS, Joo HS, Duong AC, Dieringer TD, Tan VY, Song Y, et. al. Essential *Staphylococcus aureus* toxin export system. Nat Med. 2013; 19(3):364–7.

31. Joo HS, Chatterjee SS, Villaruz AE, Dickey SW, Tan VY, Chen Y, et. al. Mechanism of Gene Regulation by a *Staphylococcus aureus* Toxin. MBio. 2016; 7(5): pii: e01579-16.

32. Cheung GY, Duong AC, Otto M. Direct and synergistic hemolysis caused by Staphylococcus phenol-soluble modulins: implications for diagnosis and pathogenesis. Microbes Infect. 2012; 14(4): 380–6.

33. Yarwood JM, Schlievert PM. Quorum sensing in Staphylococcus infections. J Clin Invest. 2003; 112(11): 1620–5.

34. Periasamy S, Joo HS, Duong AC, Bach TH, Tan VY, Chatterjee SS, et.al. How *Staphylococcus aureus* biofilms develop their characteristic structure. Proc Natl Acad Sci U S A. 2012; 109(4): 1281–6.

35. Cheung GY, Joo HS, Chatterjee SS, Otto M. Phenol-soluble modulins--critical determinants of staphylococcal virulence. FEMS Microbiol Rev. 2014; 38(4): 698–719.

36. Peschel A, Otto M. Phenol-soluble modulins and staphylococcal infection. Nat Rev Microbiol. 2013; 11(10): 667–73.

37. Cheung GYC, Fisher EL, McCausland JW, Choi J, Collins JWM, Dickey SW,et. al. Antimicrobial Peptide Resistance Mechanism Contributes to *Staphylococcus aureus* Infection. J Infect Dis. 2018; 13;217(7): 1153-1159.

38. Novick RP. Autoinduction and signal transduction in the regulation of staphylococcal virulence. Mol Microbiol. 2003; 48(6): 1429–49.

39. Grundmann H, Aires-de-Sousa M, Boyce J, Tiemersma E. Emergence and resurgence of methicillin-resistant *Staphylococcus aureus* as a public-health threat. The Lancet. 2008; 368: 874–875.

40. Tomasz A, Drugeon HB, de Lencastre HM, Jabes D, McDougall L, Bille J. New mechanism for methicillin resistance in Staphylococcus aureus: clinical isolates that lack the PBP 2a gene and contain normal penicillin-binding proteins with modified penicillin-binding capacity. 1989; 33(11): 1869-74.

41. Kato F, Kadomoto N, Iwamoto Y, Bunai K, Komatsuzawa H, Sugai M. Regulatory mechanism for exfoliative toxin production in *Staphylococcus aureus*. Infect Immun. 2011; 79(4):1660–70.

